# The processing of semantic complexity and co-speech gestures in schizophrenia: a naturalistic, multimodal fMRI study

**DOI:** 10.1101/2021.05.18.444612

**Authors:** Paulina Cuevas, Yifei He, Miriam Steines, Benjamin Straube

**Author notes:** Contributed equally (Paulina Cuevas, Yifei He). Corresponding address: Yifei He, Department of Psychiatry and Psychotherapy, Philipps University Marburg, Rudolf-Bultmann-Str. 8, 35039, Marburg, Germany, Corresponding.

## Abstract

Schizophrenia is marked by aberrant processing of complex speech and gesture, which may contribute functionally to its impaired social communication. To date, extant neuroscientific studies of schizophrenia have largely investigated dysfunctional speech and gesture in isolation, and no prior research has examined how the two communicative channels may interact in more natural contexts. Here, we tested if patients with schizophrenia show aberrant neural processing of semantically complex story segments, and if speech-associated gestures (co-speech gestures) might modulate this effect. In a functional MRI study, we presented to 34 participants (16 patients and 18 matched-controls) an ecologically-valid retelling of a continuous story, performed via speech and spontaneous gestures. We split the entire story into ten-word segments, and measured the semantic complexity for each segment with idea density, a linguistic measure that is commonly used clinically to evaluate aberrant language dysfunction at semantic level. Per segment, the presence of numbers of gestures varied (n = 0, 1, +2). Our results suggest that, in comparison to controls, patients showed reduced activation for more complex segments in the bilateral middle frontal and inferior parietal regions. Importantly, this neural aberrance was reduced in segments presented with gestures. Thus, for the first time with a naturalistic multimodal stimulation paradigm, we show that gestures reduced group differences when processing a natural story, probably by facilitating the processing of semantically complex segments of the story in schizophrenia.

## 1. Introduction

Schizophrenia is characterized by impairments in social interaction and communication, which are often conducted in a multimodal form including auditory speech and hand gestures — two of the most prominent communicative channels in daily situations (1–6). In schizophrenia, on the one hand, the production and perception of auditory speech at multiple levels of the linguistic hierarchy are impaired (7–12). On the other, it has been suggested that patients with schizophrenia suffer from a range of dysfunctions in the preparation, execution, perception, and understanding of gestures (13–19). As a result, impairments in both speech and gesture processing may directly contribute to deficits during multimodal daily communication. To date, extant neuroscientific studies of schizophrenia have primarily investigated speech and gesture impairments in isolated and strictly-controlled experimental paradigms. It remains unclear how both communication channels are processed (e.g., the observation of gesture, the comprehension of semantics) interactively by patients in an ecologically-valid and naturalistic multimodal context.

### 1.1. Schizophrenia: dysfunctional gesture processing in a naturalistic setting?

Gestures are spontaneous hand movements that deliver either repeating or complementary semantic information in addition to speech. Intact functioning of gesture has a positive impact on daily communication (20,21), but is often impaired in schizophrenia (16). In production, patients tend to produce fewer gestures (22), and they have difficulties imitating and producing pantomimes (13–15,23,24). Meanwhile, dysfunctional gesture processing has been reported. Patients showed impaired functional brain connectivity when integrating speech and gesture (25). At higher-semantic levels, their understanding of gestures is reported to be dependent on language context. For example, when processing gestures in an abstract sentence context (metaphoric gestures), patients showed impaired behavioral performance to match the semantic information of gesture to a corresponding sentence (26,27), even more so for those with more severe positive formal thought disorders (17). Of note, even if the processing of abstract gestures is impaired in schizophrenia (18,28), when the same information is presented bimodally (speech accompanied by gesture), fMRI results suggest that patients showed comparable brain activations in comparison to controls (28). These prior studies, together, may suggest that patients’ processing of gesture and its semantics may benefit from a multimodal context, as they would mostly encounter in daily naturalistic situations.

### 1.2. Idea density as a window into language deficits in schizophrenia

A naturalistic approach, which employs naturalistic and ecologically-valid experimental stimulation, has contributed significantly to a sharpened understanding of the neurobiology of language (29–32). However, this novel approach hasn’t been sufficiently exploited in clinical neuroscience research (8). Notably, clinicians have commonly used a measure called idea density (ID) to shed light on language dysfunctions in clinical populations. Idea density is a metric of semantic complexity that measures how many propositions are expressed within a specific number of words. Deriving from narratives, its ecological validity enables researchers to investigate cognitive demands and language complexity without interfering directly with the acquisition of experimental data (33). Idea density is proven informative in investigating the relationship between early linguistic abilities and in predicting the development of Alzheimer’s disease (34,35), mild cognitive impairment (36), speech impairments of patients with aphasia (37,38), and aging effects of language (39–42). More relevant to the current research, in schizophrenia, idea density has been used to understand patients’ higher-order semantic performance (33,42–44). Despite substantial insights, studies on schizophrenia to date have primarily focused on how patients *produce* narratives in either written or oral form; how patients *process* different parts of naturalistic stimuli differing in the degree of semantic complexity has not been studied. Besides, no studies have investigated the neural correlates of the processing of idea density in clinical populations, e.g., in schizophrenia. Moreover, how patients with schizophrenia, when processing a natural story, might be impaired or benefit from a *multimodal* context (i.e., processing both speech and gesture) remains unclear. In a recent study with a healthy college student sample, we examined the processing of a natural, multimodal story with fMRI, and observed a facilitative effect of co-speech gesture when the story is semantically more complex, i.e., when idea density is higher (45). Thus, it would be worthwhile to examine if patients with schizophrenia may benefit from a multimodal context in a natural context, as have been previously shown in more controlled experiments (28)

### 1.3. Present study

Thus, in the current study, we employed a naturalistic approach in an fMRI study to investigate neural correlates of dysfunctional speech and gesture processing in schizophrenia. We focused on three specific research questions for the processing of this multimodal story: 1) if the observation of gestures is impaired in schizophrenia, 2) if patients are impaired in the processing of semantic complexity of a natural story (as parameterized by idea density), and 3) if a multimodal context (with vs. without gesture) modulates semantic deficits (if there are any) in schizophrenia.

## 2. Methods

### 2.1. Participants

We summarized participants’ demographic and clinical characteristics in Table 1. Healthy controls were recruited, matching age, gender, and education to patients. Sixteen patients were recruited at the Department of Psychiatry and Psychotherapy at the Philipps University of Marburg, and were diagnosed according to ICD-10 with schizophrenia (F20.0, n=12, and F20.3, n=1) or schizoaffective disorder (F25.0, n=2, and F25.2, n=1). Participants in both groups are native speakers of German. All except one patient received antipsychotic treatment; six were additionally treated with antidepressant medication. Positive and negative symptoms were assessed with the Scale for the Assessment of Positive Symptoms (SAPS) and the Scale for the Assessment of Negative Symptoms (SANS). Eighteen age–, gender–, and education-matched healthy participants with no history of any mental disorders were recruited from the same area. Exclusion criteria for both groups were brain injury and neurological or other medical diseases affected by brain physiology. In both groups, we conducted neuropsychological tests to assess working memory function, digit span, trail making (TMT), verbal IQ (MWT-B) (46), and gesture production and perception (BAG, Brief Assessment of Gesture (47). These measures are reported in Table 1. All participants had normal or corrected-to-normal vision and hearing. Except for one control and one patient, all other participants are right-handed (Oldfield, 1971). All participants gave written informed consent prior to participation in the experiment and were compensated monetarily. The study was approved by the ethics committee of the School of Medicine, Philipps University Marburg.

**Table 1.** Demographic, medication, symptom, and neuropsychological measures.

|  | Patients (n=16) | Controls (n=18) |
| --- | --- | --- |
| Age (years) | 34 (12.18) | 31.94 (10.21) |
| Gender male/female | 13/3 | 13/5 |
| Education (years) | 11.81 (1.83) | 12.72 (1.36) |
| TMT A (seconds) | 31.47 (11.08) | 26.17 (9.89) |
| TMT B (seconds) | 69.30 (39.01) | 52.93 (19.58) |
| Digit Span forward | 7.75 (1.61) | 8.05 (2.43) |
| Digit Span backward | 6.00 (1.31) | 6.61 (2.50) |
| Verbal IQ | 28.87 (5.42) | 28.5 (3.79) |
| BAG subscores | 3.20 (0.53) | 3.37 (0.46) |
| SAPS (global) | 15 (6.89) |  |
| SANS (global) | 9 (6.02) |  |
| CPZ Equivalent | 562.52 (372.63) |  |

Values are presented as mean (SD). Pairwise t-tests for all reported measures were non-significant (all p-values > 0.12). TMT: trail making test; CPZ: chlorpromazine.

### 2.2. Stimuli

All participants were presented with 16 video clips depicting an actor (Figure 1) narrating consecutive parts of a slightly modified version of the short story “Der Kuli Kimgun” (48). The story was unfamiliar to all participants. The trained actor narrated the story naturally and performed spontaneous gestures of any kind (iconic, metaphoric, beat, and emblematic) using hands and arms. The actor decided freely the moment and the way to make the gestures, which were all congruent with the semantic content of the story. The presentation of the videos lasted 32:12 minutes, with individual clips lasting between 1:02 and 3:31 minutes (for detailed information, see (45,49)).

**Figure 1:**
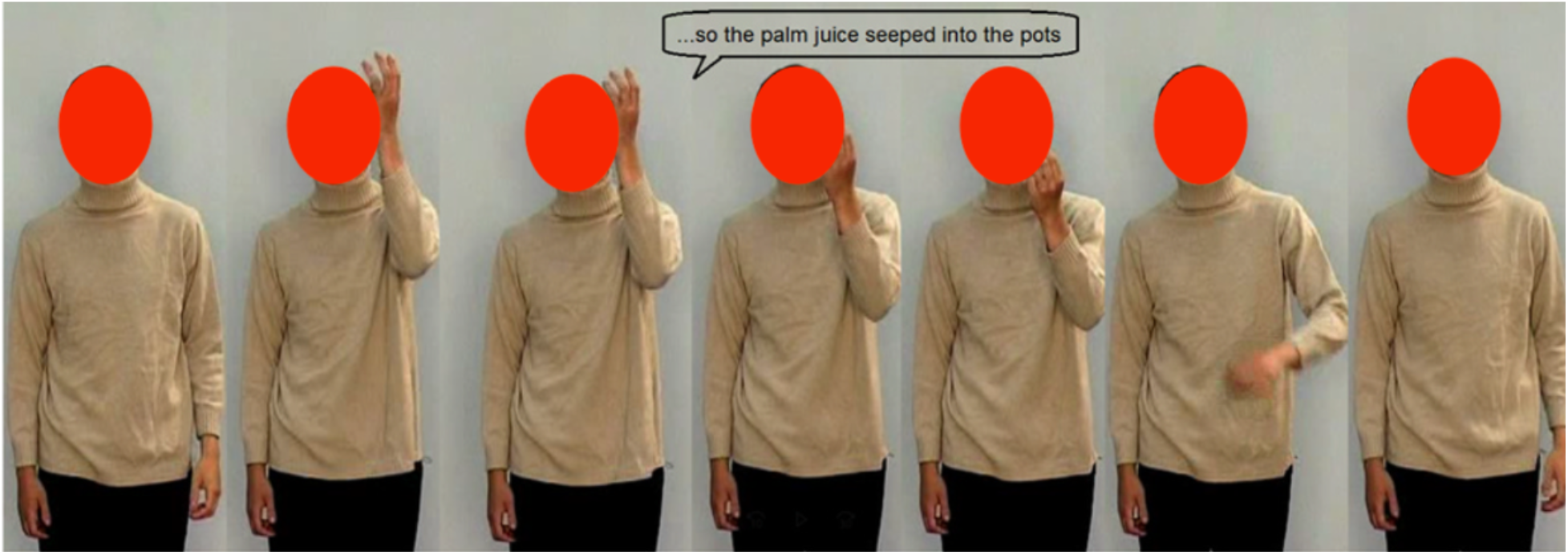
Example of stimuli. Seven still frames of the video illustrate how the actor performed the gestures (45). A short segment of the story was translated into English and depicted in the speech bubble for illustrative purposes. The actor moved his fingers in the air while moving down his arm, indicating how the liquid (palm juice) ran down into the pots. Note that in this preprint the face area of the actor is masked. In the experiment, the face of the actor remains visible for all participants.

### 2.3. Experimental design and procedure

The entire story was divided into 330 segments of 10 words each, in order to identify parts of the story with low or high semantic complexity. As there were jitter periods of 6-14 seconds between videos, we thus excluded text segments that included the end of one video and the beginning of the next one from the analysis. Then, the semantic complexity of each remaining 10-word-segment was measured by the ID (for more details about the calculation, see (45)). Every segment of the story had an ID value between two and nine (mean: 5.35 SD: 1.12). The segments with an ID value of five or below were counted as low complexity segments and segments with a value of six or higher were considered as high complexity ones. In addition, each segment included between zero and four gestures (mean: 1.50, SD: 0.82). The number of gestures appearing in each segment did not differ significantly between the low- and high-ID conditions (see (45)). Gestures that occurred between two segments were counted twice: once in the segment where they started, and once in the segment where they ended. Based on the ID value (low vs. high) and the number of gestures per segment, six different conditions were defined: zero gestures (low_noG, high_noG), one gesture (low_1G, high_1G), and two or more gestures (low_2+G, high_2+G). No significant interaction for the number of trials and segment duration was found between the factors *gesture* and *ID*. However, we had fewer trials without gestures and longer duration for the high complexity conditions. This was regarded as unproblematic as our analyses focused on the interaction between both factors (for the complete description of the stimulus material, see (45).

During the acquisition of fMRI data, participants were asked to attend to the videos (presented through a mirror mounted on the head coil) and to watch and listen to the actor carefully. Air conduction headphones (Siemens) were used for the presentation of verbal information. The loudness of the headphones was kept constant across sessions. In order to make sure that the subjects listened to the narrative, they were asked to answer a questionnaire containing details of the story (e.g., critical moments, narrative shifts, etc.).

### 2.4. fMRI data acquisition

All images were acquired using a 3-Tesla Siemens scanner (Siemens MRT Trio series). The functional images were obtained using a T2*-weighted echo-planar image sequence (TR = 2 s, TE = 30 ms, flip angle = 90°, slice thickness = 4 mm, interslice gap = 0.36 mm, field of view = 230 mm, matrix = 64 × 64, voxel size = 3.6 × 3.6 × 4.0 mm, 30 axial slices orientated parallel to the AC-PC line). 970 functional images were acquired (450 during the first run and 520 during the second run). The first acquisition run lasted 15 minutes and the second one lasted 17 minutes and 20 seconds, due to the varying lengths of individual videos. Simultaneously to fMRI data, additional EEG data were acquired. These data were intended to be used for another research project and will not be further discussed here (50,51). The use of an EEG cap was not expected to affect the quality of the BOLD responses (52).

### 2.5. Data analysis

The MRI images were analyzed using Statistical Parametric Mapping (SPM12; www.fil.ion.ucl.uk) implemented in MATLAB 2009b (Mathworks Inc. Shevorn, MA). To minimize T1-saturation effects, the first two volumes were discarded from the analysis. Afterward, all images were registered to the first image of the first run and co-registered to the anatomical volume, normalized into MNI space (voxel size = 2 × 2 × 2 mm), and smoothed with an 8 mm isotropic Gaussian filter. A high-pass filter (cut-off period 128 sec) was used. Statistical analysis was performed in a two-level procedure. The design matrix for the modulation of single-subject BOLD responses at the first level comprised the onsets and durations of all six conditions, as well as the six movement parameters of each subject. The hemodynamic response function (HRF) was modeled by the canonical HRF. A flexible factorial second-level analysis with six conditions (low_noG, low_1G, low_2+G, high_noG, high_1G, high_2+G) was performed.

A Monte Carlo simulation of the brain volume of the current study was employed to determine an adequate voxel contiguity threshold (53). It is suggested that this correction provides sensitivity to smaller effect sizes and also corrects for multiple comparisons across the whole brain volume (54). Assuming an individual voxel type 1 error of p < .001, a cluster extent of 87 contiguous resampled voxels was indicated as necessary to correct for multiple comparisons at p < .05. Thus, clusters with at least 87 voxels and a significance level of p < .001 are reported for all contrasts. All described coordinates of activation are located in MNI space. The AAL toolbox (55) was employed for the anatomical localization of the clusters.

### 2.6. Contrasts of interests

We firstly tested for general group differences in the neural processing of the narrative and group differences in the processing of gestures (group × gesture). We also tested, across groups, if gesture facilitates the processing of story differing in complexity, as have been observed in (45,49). Then, we focused on the between-group difference in the processing of story segments differing in ID (group × complexity). Lastly, we tested the interaction of group × complexity × gestures to test if group-specific processing of semantic complexity may be modulated by the presence of co-speech gestures.

## 3. Results

We found no significant main effect of group and no group differences regarding the processing of gestures. Post-hoc tests revealed an activation increase in both groups during the processing of co-speech gestures (*Speech + Gesture [gesture:1&2+] > Speech [gesture:0]*) in the bilateral temporo-occipital regions (Figure 2B, 2D and Table 1s). The activation in this region increased according to the number of gestures presented, similarly across both groups, as suggested by the results from the conjunction analysis between controls and patients (Figure 1s, Table 3s, supplement).

**Figure 2:**
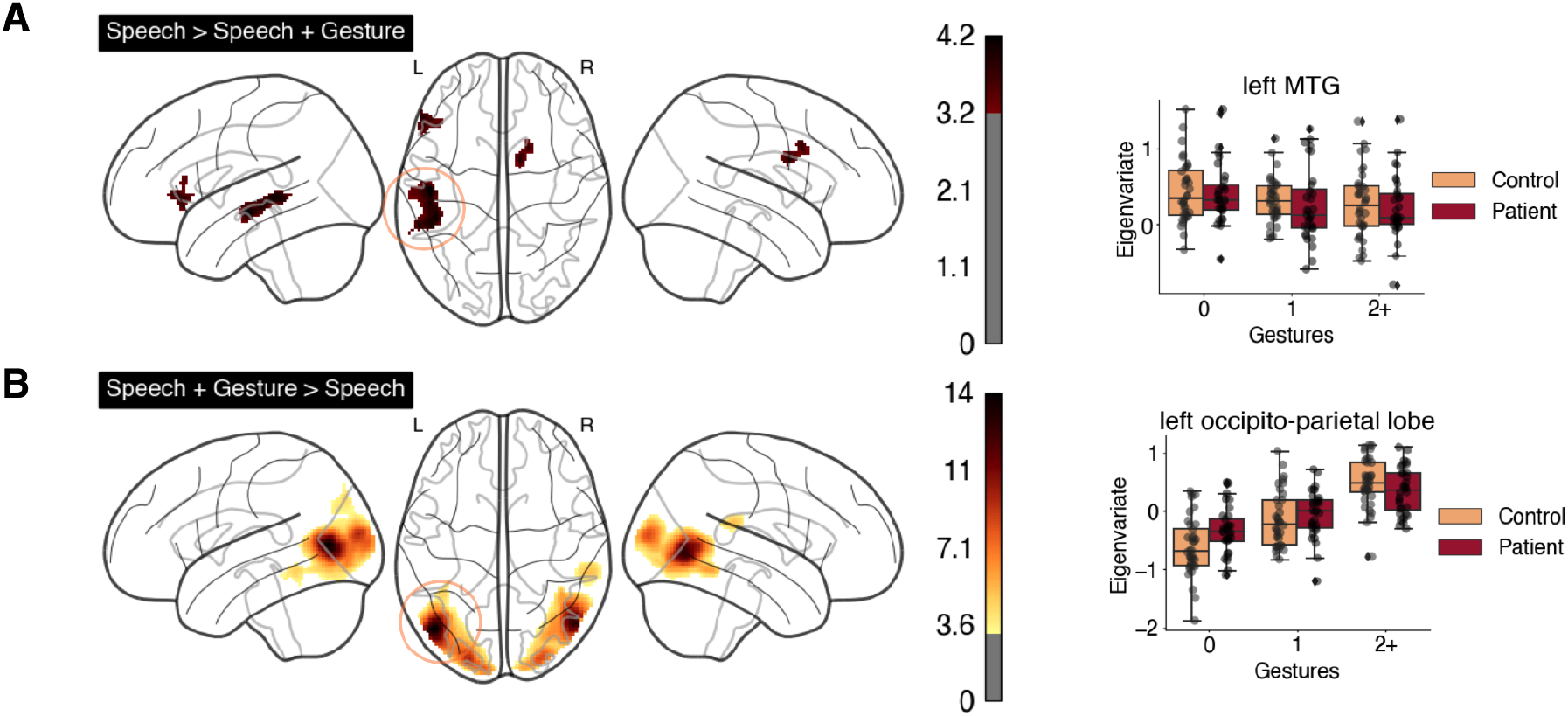
Gesture-related activation decrease and activation increase in both groups. (A) Contrast of [Speech > Speech + Gesture] as shown in the glass brain (left panel) and boxplots (right panel) showing eigenvariates extracted from the cluster that peaks at the left middle temporal gyrus (left MTG, [−48, −16, −10; k = 444]). (B) Contrast of [Speech + Gesture > Speech] as shown in the glass brain (left panel) and boxplots (right panel) showing eigenvariates extracted from the cluster that peaks at the left occipital lobe [−46, −72, −6; k = 4092]. For all glass brain figures, the threshold for voxel activations was set at p < 0.001 uncorrected, and only clusters larger than 87 voxels have been included (Monte-Carlo cluster-extent corrected at p < .05). Color bar indicates the scale of the T-statistics. Speech: Gesture = 0; Speech + Gesture: Gesture = 1 & 2+.

The reverse contrast (*Speech [gesture:0] > Speech + Gesture [gesture:1&2+]*) indicates that gestures resulted in group-independent reduced activation of the left middle temporal gyrus (MTG), the left inferior frontal gyrus (IFG), and the right caudate (see Figure 2A, 2C and Table 1s in the supplement). Again, conjunction analysis suggests that left MTG activation is comparable across groups (Figure 1s, Table 3s, supplement).

We report group-independent interaction analysis of *gesture × complexity* in the supplement, Table 2s, and Figure 2s. The findings replicated results from our previous studies (45,49), suggesting that gesture reduced the activation in the left temporal lobe only when semantic complexity is high (high-ID).

The contrast *group × complexity* revealed that, in comparison to controls, patients showed reduced activation in the left inferior parietal lobe (left IPL) when ID is high; however, in the low ID passages, the left IPL activation in patients was higher than that of controls (Figure 3 and Table 1s).

**Figure 3:**
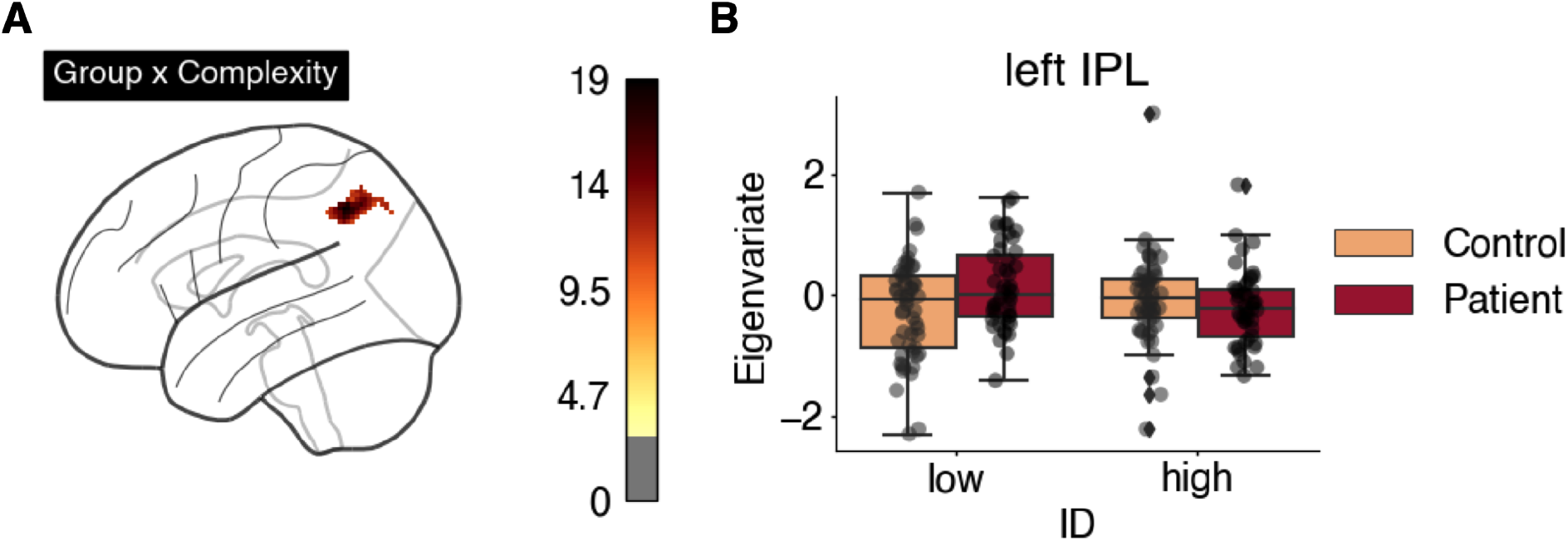
(A) Interaction *group × complexity* shown in the glass brain figure. (B) Boxplots showing eigenvariates extracted from the cluster that peaks at the left inferior parietal lobe (left IPL, [−52, −54, 44; k = 149]). The threshold for the glass brain figure was set at p < 0.001 uncorrected, and only clusters larger than 87 voxels have been included (Monte-Carlo cluster-extent corrected at p < .05). Color bar indicates the scale of the F-statistics.

Lastly, the interaction *group × complexity × gesture* (Figure 4, Table 1s) activated the bilateral IPL and the middle frontal gyrus (MFG), and additionally the right IFG. Here, in these regions, the activation increase for higher ID passages (high-ID > low-ID) was observed only when no gestures were presented (Gestures = 0). When the story was presented in a multimodal form (Gestures = 1 & 2+), this high-ID > low-ID difference was normalized for patients.

**Figure 4:**
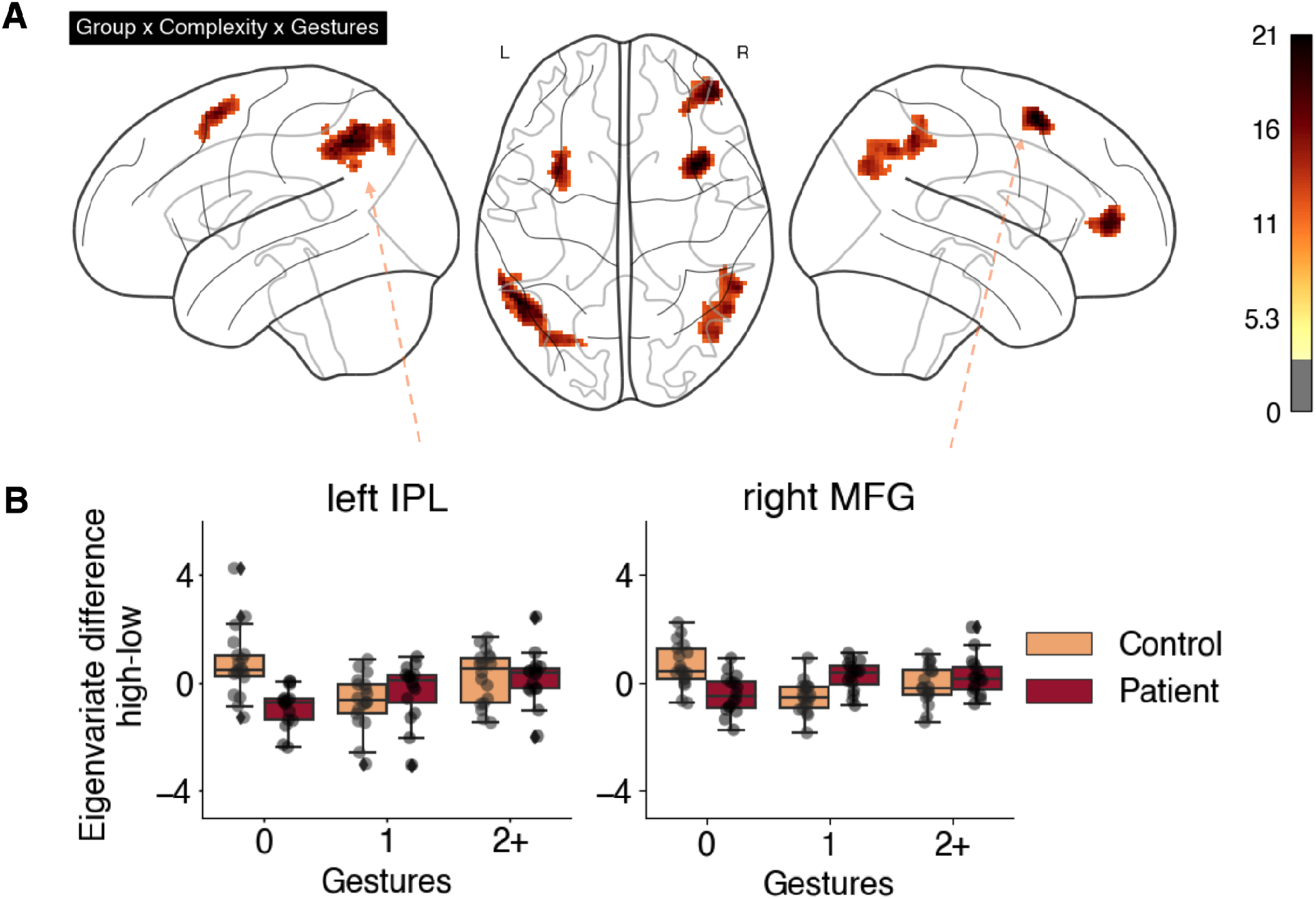
(A) Interaction *group × complexity × gestures* shown in the glass brain figure. (B) Boxplots showing the difference of eigenvariates between high-ID and low-ID conditions, extracted from the clusters that peak at the left inferior parietal lobe (left IPL, [−50, −56, 46; k = 367]), and the right middle frontal gyrus (right MTG [−34, 12, 58; k = 181]). The threshold for the glass brain figure was set at p < 0.001 uncorrected, and only clusters larger than 87 voxels have been included (Monte-Carlo cluster-extent corrected at p < .05). Color bar indicates the scale of the F-statistics.

## 4. Discussion

In the current study, we investigated if patients with schizophrenia are impaired in the neural processing of semantic complexity within a naturalistic multimodal context. Our results suggest that patients, in comparison to controls, showed increased activation of the left IPL during comprehending semantically less complex passages; however, the left IPL activation was reduced for patients when semantic complexity is high. More importantly, the presentation of gestures reduced this group difference, suggesting a facilitative or compensatory role of gestures in this context.

### 4.1. The processing of gesture is comparable between groups

For both groups, the *Speech + Gesture > Speech* comparison activated the bilateral posterior temporo-occipito-parietal visual regions, thus suggesting that patients are intact in recruiting this region to process gesture when processing naturalistic stimuli, in a similar way to isolated bimodal videos (27,45,51,56). These findings may imply that, despite well-documented gesture dysfunctions in schizophrenia (13,57), when engaged in a high-level context (i.e., presented together with speech in both isolated and naturalistic multimodal settings), the perception of gestures in schizophrenia could appear to be largely intact.

Results from the *Speech > Speech + Gesture* comparison are in accordance with our prior studies (45,49), and suggest that segments without gestures may recruit more neural resources in the left hemispheric speech and language regions (e.g., the left IFG and MTG. Importantly, this facilitative effect of gesture was present for both patients and controls, thus suggesting that gesture may facilitate story comprehension in general in schizophrenia.

### 4.2. Aberrant neural processing of semantic complexity in schizophrenia is modulated by gesture

The focus of this study was on potential impairments when patients with schizophrenia process the naturalistic multimodal story differing in the degree of semantic complexity. Indeed, when collapsing across all gesture conditions (gesture: 0, 1, 2+), patients, as opposed to controls, showed increased activation for the low complexity and decreased activation for high complexity passages in the left IPL. Moreover, when the factor of gesture is considered, we observed that this effect interacts with gesture in the bilateral IPL and additionally the middle frontal regions. Specifically, the neural aberrance in these regions (including the left IPL) was only present when no gesture was presented. In contrast, when the story was presented in a multimodal form (gesture: 1 and 2+), group differences were largely not observable.

The left IPL is an important region that underlies a range of cognitive functions that are functionally relevant to human social communication (58,59). Importantly, the left IPL is crucially involved during language comprehension at the semantic level (60,61), and has been reported to support semantic prediction during comprehension of a naturalistic auditory story (31). Besides, the left IPL is associated with working memory and is sensitive to the cognitive load of the task (62). In schizophrenia, the left IPL activity is reported to be reduced for patients when they are engaged in sentence-level language comprehension tasks (63). In the current study, although patients showed even increased activation in the left IPL in the low-complexity passages, when they process parts of the story that are semantically more complex, they failed to activate the left IPL as controls did. This neural aberrance, during the processing of a naturalistic narrative, might derive from patients’ impaired working memory capacity and semantic processing (7,64–66).

The left middle frontal gyrus (MFG) is associated with the maintenance of verbal information (67). In this regard, participants might recruit this region especially during the complex speech conditions because multiple semantic events have to be kept in mind in order to interpret the entire story. Nevertheless, patients recruit this region already even to a higher degree than controls, during the low complexity speech condition. However, in the high complexity condition, the left MFG activity in patients was reduced. Notably, a study investigating the processing of novel and conventional metaphors by patients with schizophrenia also reported increased activation of the left MFG (68), importantly, there, the activation decreased as the cognitive demands of the task increased (in this case abstract thinking). Similarly, in our study, patients’ activation in this region decreased as the demands of the narrative increased, i.e., in high complex passages. On the other hand, the right MFG plays an important role in the temporal sequencing of discourse (69), an ability that is fundamental for interpreting the story.

Together, for the bilateral IPL and MFG, the increased activation by patients during the low complexity condition indicate that patients already need more neural effort (increased BOLD response) for the processing of low complexity segments; when the story is semantically too complicated (high-ID), patients then failed to provide the necessary resources in these regions (reduced BOLD response), partially due to their impaired working memory maintenance or processing of complex semantics.

However importantly, this neural aberrance was only observable when there was no gesture presented together with speech. In contrast, in these regions, whenever the story was presented in a multimodal form (gesture: 1 and 2+), the activation pattern is comparable between groups. A similar facilitative effect of gesture when processing a story has been observed for elder vs. younger participants, i.e., group difference was reduced when gestures were presented together with story segments (49). In line with these findings, for isolated sentences, despite the fact that patients showed neural aberrance when processing abstract-social gestures and concrete-non-social speech, group difference was significantly reduced when these events were presented in a multimodal form (28). In the current study, we extend prior findings by showing a similar facilitative nature of gesture during speech comprehension: In a naturalistic story setting, even if patients neural aberrance may be more likely to derive from their impaired semantic prediction and working memory capacity, which are not as heavily demanded in isolated situations, nevertheless, a multimodal context is, again, proven beneficial (6,21).

### 4.3 Limitation

Despite new insights, the current study has clear limitations. Limited by the sample size of this study (n = 16 & 18 for each group), we are unable to test the role of medication for story comprehension, and cannot pinpoint if the observed neural aberrance (as well as neural facilitation) is specific to chronic or first-episode schizophrenia. Therefore, future studies are expected to investigate language processing and gesture–speech interaction in schizophrenia with naturalistic stimulation with larger samples.

## 5. Conclusion and future research

The following conclusions can be drawn from this naturalistic experiment: First, patients and controls recruit similar regions for the processing of gestures and speech in general. Second, patients, when compared to controls, show reduced activation in the bilateral middle frontal and parietal regions when processing parts of the story that are semantically more complex. Finally, gestures reduced group differences, potentially by facilitating the processing of the narrative. Thus, future studies could examine if the differences in the processing of semantic complexity are reduced by conducting existing training developed to improve verbal communication and social cognition (70,71)

## Supporting information

suppplement

## 6. Acknowledgments

This work was supported by the International Research Training Group ‘‘The Brain in Action’’ (IRTG 1901) by the German Research Foundation (DFG), the von Behring-Röntgen-Stiftung (vBR59-0002, 64-0001), the Hessisches Ministerium für Wissenschaft und Kunst (HMWK; project ‘The Adaptive Mind’), and the individual projects of the DFG (HE8029/2-1). The study was also supported by the Core Facility Brain Imaging, Faculty of Medicine, University of Marburg, Rudolf-Bultmann-Str. 9, 35039, Marburg, Germany.

