## Supplementary material for "The processing of semantic complexity and co-speech gestures in schizophrenia: a naturalistic, multimodal fMRI study": suppplement

**A naturalistic and multimodal fMRI study’**

***Table 1s****: Activation peaks and cluster extents*

| **Contrast** | **Anatomical region** | **Cluster extent** | **Hem.** | **MNI Coordinates**  **x y z** | | | ***T*-value** | **No. voxels** |
| --- | --- | --- | --- | --- | --- | --- | --- | --- |
| **SSD >Controls** | | | | | | | | |
|  |  |  |  |  | | | ***n.s.*** |  |
| **Controls > SSD** | | | | | | | | |
|  |  |  |  |  | | | ***n.s.*** |  |
| **High complexity > low complexity** | | | | | | |  |  |
| Cerebellum | | Vermis | R | 6 | -68 | -30 | 4.34 | 197 |
| **Low complexity > high complexity** | | | | | | | | |
|  | |  |  |  |  |  | **n.s.** |  |
| **Gesture+speech > Speech** | | |  |  |  |  |  |  |
| Middle occipital gyrus | | Middle temporal gyrus, inferior occipital gyrus, superior occipital gyrus, cuneus, fusiform gyrus, inferior temporal gyrus | L | -46 | -72 | 6 | 14.25 | 4092 |
| Middle temporal gyrus | | Middle occipital gyrus, inferior temporal gyrus, superior occipital gyrus, cuneus, fusiform gyrus, middle temporal gyrus | R | 48 | -68 | 2 | 12.82 | 3184 |
| Superior temporal gyrus | | Superior temporal, supramarginal gyrus, rolandic operculum | R | 60 | -34 | 22 | 5.06 | 237 |
| **Speech> Gesture+Speech** | | | | | | | | |
| Middle temporal gyrus | | Superior temporal gyrus, inferior temporal gyrus, hippocampus, insula | L | -48 | -16 | -10 | 4.21 | 444 |
| Caudate | | Anterior cingulate, middle cingulate, superior frontal gyrus, middle frontal gyrus | R | 12 | 4 | 26 | 3.87 | 97 |
| Pars triangularis | | Pars orbitalis, middle frontal gyrus, Insula | L | -54 | 32 | 2 | 3.83 | 149 |
| **Interaction gesture x group** | | | | | | | **F-value** |  |
|  | | | | | | | **n.s.** |  |
| **Interaction complexity x group** | | | | | | | **F-value** |  |
| Inferior parietal gyrus | | Angular gyrus, supramarginal gyrus, superior parietal gyrus | L | -52 | -54 | 44 | 18.99 | 149 |
| **Interaction gesture x complexity x group** | | | | | | | **F-value** |  |
| Middle frontal gyrus | | Superior frontal gyrus, precentral gyrus | R | 34 | 12 | 58 | 21.39 | 181 |
| Inferior parietal gyrus | | Inferior parietal gyrus, angular gyrus, supramarginal gyrus, superior parietal gyrus | L | -50 | -56 | 46 | 21.34 | 367 |
| Middle frontal gyrus | | Pars triangularis, pars orbitalis, insula, superior frontal gyrus | R | 44 | 46 | 4 | 19.11 | 231 |
| Middle frontal gyrus | | Superior frontal gyrus, precentral gyrus | L | -32 | 10 | 60 | 17.66 | 154 |
| Inferior parietal gyrus | | Supramarginal gyrus, superior parietal gyrus, angular gyrus, middle occipital gyrus | R | 56 | -48 | 48 | 17.12 | 375 |

Voxel activations thresholded at p < 0.001, only clusters bigger than 87 voxels are reported (Monte-Carlo cluster corrected at p < 0.05). Lateralization of the activation clusters is indicated by L (left) and R (right).

**Table 2s**: Activation peaks and cluster extents of the group-independent interaction gesture x ID

| **Contrast** | **Anatomical region** | **Cluster extent** | **Hem.** | **MNI Coordinates**  **x y z** | | | ***F*-value** | **No. voxels** |
| --- | --- | --- | --- | --- | --- | --- | --- | --- |
| Interaction gesture x ID | | | | | | | | |
| Superior temporal pole | | Insula, pars orbitalis, middle temporal pole, middle temporal gyrus | L | -46 | 12 | -12 | 20.90 | 287 |
| Inferior occipital gyrus | | Middle occipital gyrus, fusiform gyrus | R | 28 | -90 | -2 | 19.53 | 383 |
| Cerebelum | | Vermis | R | 6 | -66 | -30 | 19.04 | 589 |
| Herschl gyrus | | Hippocampus, insula, thalamus, putamen | L | -30 | -30 | 4 | 18.61 | 199 |

Voxel activations thresholded at p < 0.001, only clusters bigger than 87 voxels are reported (Monte-Carlo cluster corrected at p < 0.05). Lateralization of the activation clusters is indicated by L (left) and R (right).

**Table 3s**: Activation peaks and cluster extents of the conjunction analysis for modality effects

|  | **Anatomical region** | **Cluster extent** | **Hem.** | **MNI Coordinates** | | | ***T*-value** | **No. voxels** |
| --- | --- | --- | --- | --- | --- | --- | --- | --- |
|  |  |  |  | **x y z** | | |  |  |
| Conjunction Control and Patients [Speech > Speech + Gesture], p<0.05, k = 100 | | | | | | | | |
| Middle temporal gyrus | | Middle temporal gyrus | L | -48 | -40 | 0 | 2.84 | 647 |
| Conjunction Control and Patients [Speech + Gesture > Speech] | | | | | | | | |
| Middle temporal gyrus | | Middle temporal gyrus, middle occipital gyrus | L | -44 | -70 | 8 | 9.06 | 1522 |
| Middle temporal gyrus | | Middle temporal gyrus, inferior temporal gyrus, inferior occipital gyrus | R | 48 | -60 | 0 | 7.54 | 917 |
| Superior occipital gyrus | | Superior occipital gyrus | L | 26 | -92 | 14 | 4.80 | 164 |

Voxel activations thresholded at p < 0.001, only clusters bigger than 87 voxels are reported (Monte-Carlo cluster corrected at p < 0.05). Lateralization of the activation clusters is indicated by L (left) and R (right).

*Figure 1S*

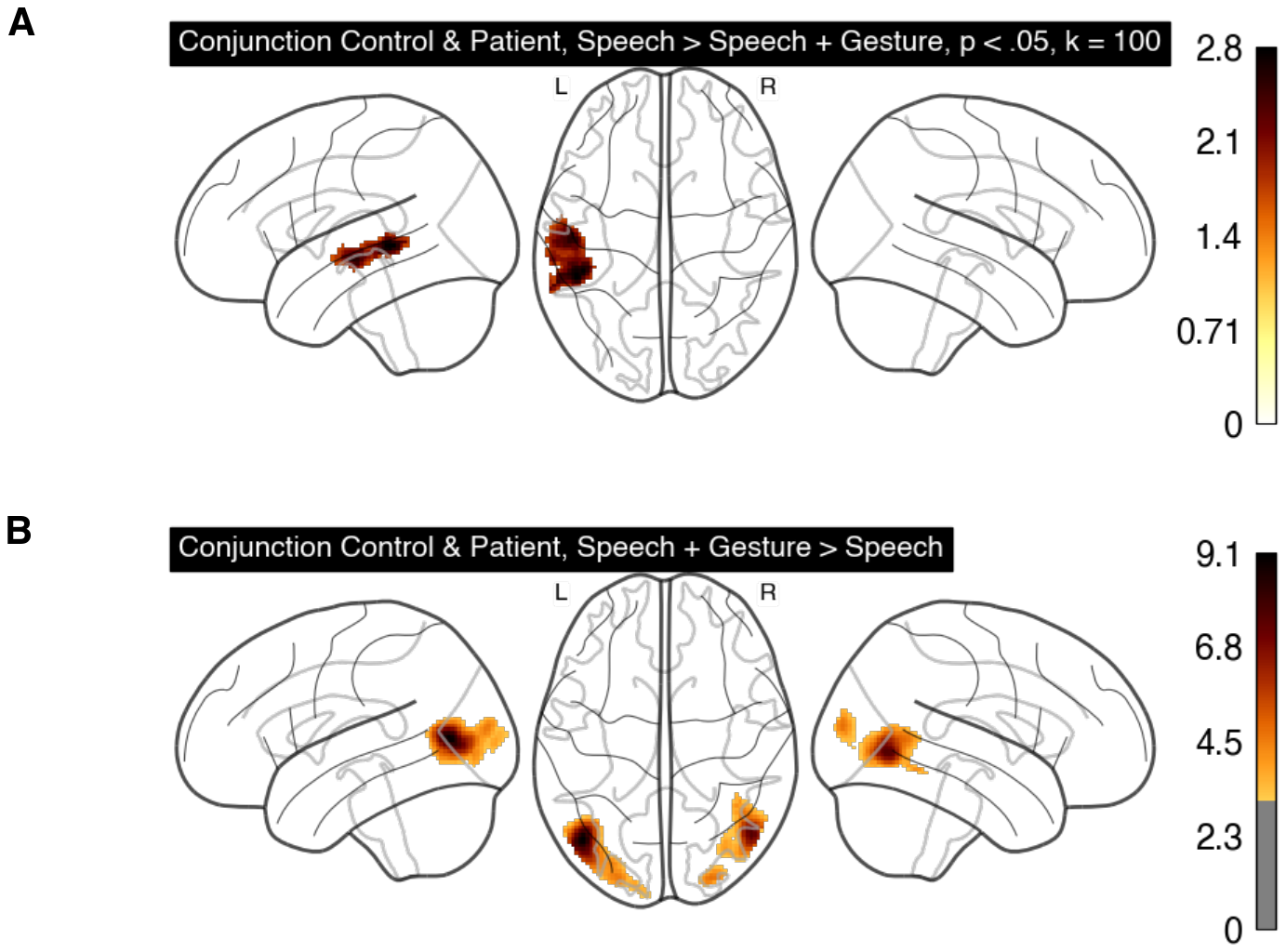

**Figure 1S:** Results from the conjunction analysis showing comparable modality effects between groups. **(**A) Conjunction between controls and patients for the contrast of [Speech > Speech + Gesture] as shown in the glass brain. Note that here the threshold for voxel activations was set at p < 0.05, k = 100, uncorrected. (B) Conjunction between controls and patients for the contrast of [Speech + Gesture > Speech]. The threshold for voxel activations was set at p < 0.001 uncorrected, and only clusters larger than 87 voxels have been included (Monte-Carlo cluster-extent corrected at p < .05). For both panels, color bar indicates the scale of the T-statistics. Speech: Gesture = 0; Speech + Gesture: Gesture = 1 & 2+).

*Figure 2S*

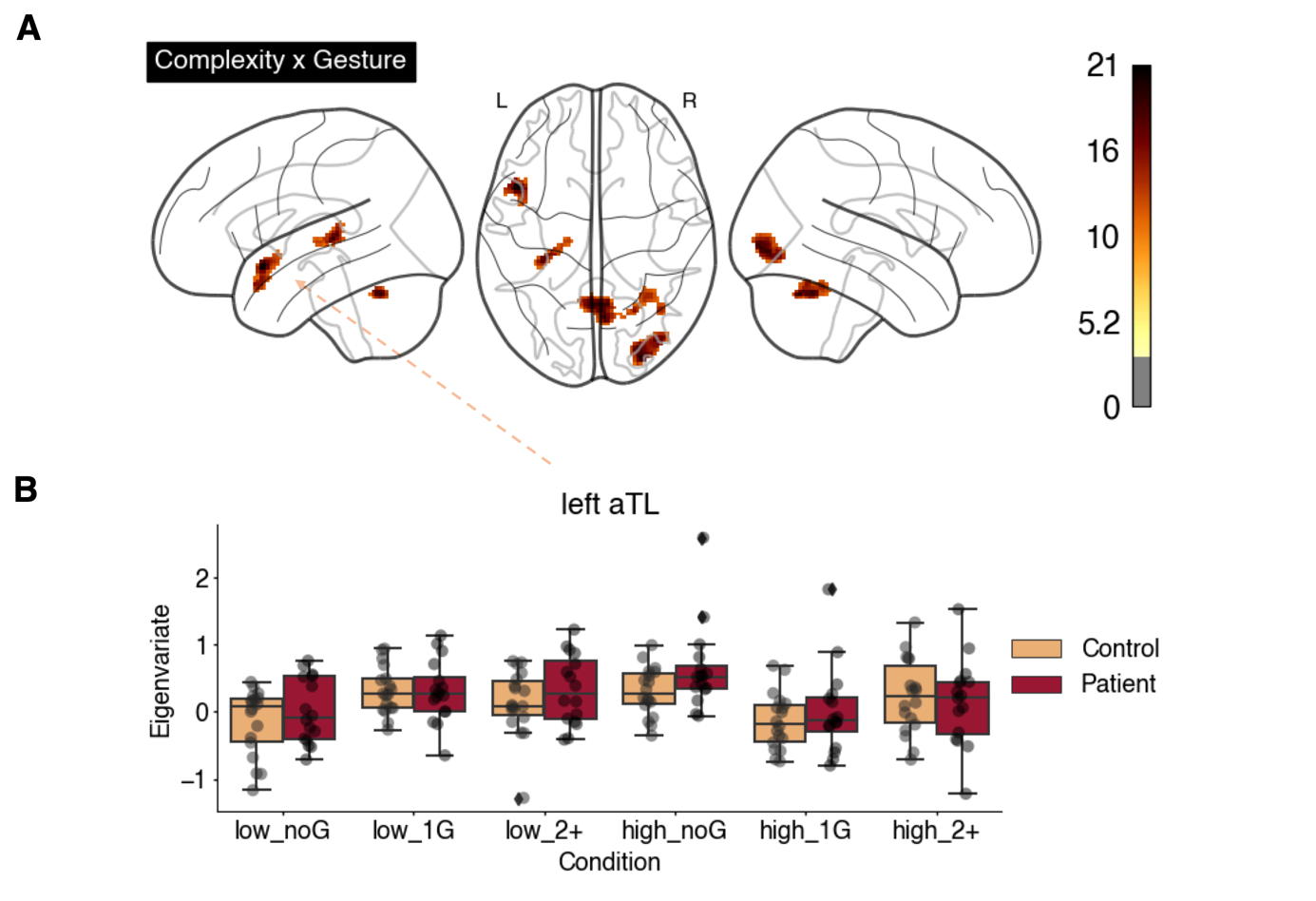

**Figure 2S:** Interaction gestures x ID**. (**A) Activation clusters localized in the anterior temporal pole, inferior occipital gyrus, cerebellum and Herschel gyrus. (B) Boxplots showing reduction of activation at the anterior temporal pole during the high complexity conditions when gesture was presented [high_1G, high_2+]. The threshold for the glass brain figure was set at p < 0.001 uncorrected, and only clusters larger than 87 voxels have been included (Monte-Carlo cluster-extent corrected at p < .05).
